# The *ivory* lncRNA regulates seasonal color patterns in buckeye butterflies

**DOI:** 10.1101/2024.02.09.579733

**Authors:** Richard A. Fandino, Noah K. Brady, Martik Chatterjee, Jeanne M. C. McDonald, Luca Livraghi, Karin R. L. van der Burg, Anyi Mazo-Vargas, Eirene Markenscoff-Papadimitriou, Robert D. Reed

**Affiliations:** Department of Ecology and Evolutionary Biology, Cornell University; Ithaca, New York, United States of America; Department of Biological Sciences, The George Washington University; Washington D.C., United States of America; Department of Biological Sciences, Clemson University; Clemson, South Carolina, United States of America; Department of Molecular Biology and Genetics, Cornell University; Ithaca, New York, United States of America

**Keywords:** Butterfly wing patterns, evo-devo, long non-coding RNA, cortex, plasticity

## Abstract

Long non-coding RNAs (lncRNAs) are transcribed elements increasingly recognized for their roles in regulating gene expression. Thus far, however, we have little understanding of how lncRNAs contribute to evolution and adaptation. Here we show that a conserved lncRNA, *ivory*, is an important color patterning gene in the buckeye butterfly *Junonia coenia*. *ivory* overlaps with *cortex*, a locus linked to multiple cases of crypsis and mimicry in Lepidoptera. Along with a companion paper by Livraghi et. al., we argue that *ivory*, not *cortex*, is the color pattern gene of interest at this locus. In *J. coenia* a cluster of *cis*-regulatory elements (CREs) in the first intron of *ivory* are genetically associated with natural variation in seasonal color pattern plasticity, and targeted deletions of these CREs phenocopy seasonal phenotypes. Deletions of different *ivory* CREs produce other distinct phenotypes as well, including loss of melanic eyespot rings, and positive and negative changes in overall wing pigmentation. We show that the color pattern transcription factors Spineless, Bric-a-brac, and Ftz-f1 bind to the *ivory* promoter during wing pattern development, suggesting that they directly regulate *ivory*. This case study demonstrates how *cis*-regulation of a single non-coding RNA can exert diverse and nuanced effects on the evolution and development of color patterns, including modulating seasonally plastic color patterns.

**Significance:** The genomic locus hosting the *cortex* gene has been linked to numerous cases of color pattern adaptation in moths and butterflies, including crypsis, mimicry, and seasonal polyphenism. Here we show in buckeye butterflies that the actual color pattern gene at the *cortex* locus is an evolutionarily conserved long non-coding RNA (lncRNA), dubbed *ivory*, that overlaps with *cortex*. Compared with other wing pattern genes, *ivory* stands out because of the highly nuanced, quantitative changes in pigmentation that can be achieved by manipulating adjacent *cis*-regulatory sequences. This study highlights how lncRNAs can be important factors underlying morphological evolution, and emphasizes the importance of considering non-coding transcripts in comparative genomics.

## Introduction

Seasonal cycles are deeply ingrained in the genomes of many animals. Evolutionary adaptation to annual fluctuations in conditions such as daylength, temperature, and precipitation is often realized in the form of phenotypic plasticity, where trait expression is determined in response to external cues. While the physiological and endocrine mechanisms underlying plasticity have been well characterized in many systems, there are still relatively few case studies that highlight the genetic basis of plasticity in natural populations. Buckeye butterflies, *Junonia coenia*, have been a useful model in this regard. Across much of their range, these butterflies are polyphenic, and show distinct seasonal color patterns – in the summer they are a light tan color, while in the autumn they are dark red and thus absorb heat more efficiently (1). This polyphenism is mediated by seasonal changes in daylength and temperature, and natural genetic variation in this response has been mapped to three loci: *herfst*, *trehalase*, and *cortex* (2). Of these, the *cortex* locus is of particular interest because it has also been implicated in adaptive melanism in moths (3), leaf mimicry in *Kallima* (4), and color pattern mimicry in *Heliconius* and *Papilio* (5-7). It remains a question of fundamental interest how a single locus like this can control such a wide variety of adaptive variation between species, populations, and even seasons. Such multifunctionality is presumably driven by *cis*-regulatory innovation, but we still have a poor understanding of how adaptive changes, including plasticity, are distributed across *cis*-regulatory regions of individual genes. Genetic association mapping coupled with chromatin accessibility comparisons in *J. coenia* have linked several *cis*-regulatory elements (CREs) at the *cortex* locus to plasticity, which now motivates us to functionally characterize how adaptive color pattern variation is encoded at this locus (2).

While there has been significant interest in the *cortex* locus for the reasons described above, functional studies have, in fact, proven vexing. First, expression work looking at *cortex* mRNA and protein localization in developing butterfly wings has shown inconsistent color pattern associations (5, 6). Second, mosaic deletion phenotypes occur at such low frequencies they could possibly be explained by spurious long deletions affecting adjacent genes or enhancers (8). And last, and most importantly, Cas9-induced germ line deletions in *cortex* show no phenotypes in F1 individuals (9). Remarkably, as we show below, and is also shown in a companion paper by Livraghi et al. (9), this all seems to be explained by the fact that *cortex* is not the color pattern gene we once thought it was – rather, the actual gene of interest is a long non-coding RNA (lncRNA), dubbed *ivory*, that partially overlaps with the *cortex* gene on the antisense strand. This is interesting because few lncRNAs have thus far been implicated in adaptation, even though lncRNAs are of major emerging interest in developmental biology due to their roles as chromatin and spatially restricted transcription regulators (10, 11).

The goals of this study, then, were threefold. First, we sought to provide functional validation of the *ivory* lncRNA as a color patterning gene in the buckeye butterfly *J. coenia*, with a focus on seasonally plastic wing pattern phenotypes (12, 13). Second, we aimed to characterize some of the *cis*-regulatory architecture of the *ivory* locus using both comparative and functional approaches. Last, using a deletion line, we aimed to assess the effects of *ivory* on gene expression during wing development. We show that the *ivory* lncRNA is a fundamental regulator of wing pigmentation with a deeply conserved *cis*-regulatory region. Deletions at different regulatory regions of the *ivory* locus not only phenocopy seasonal color pattern phenotypes but can have broader positive and negative quantitative effects on pigmentation. This work, accompanied by Livraghi et al.’s comparative study (9), calls for a major revision of our models of butterfly wing pattern development, and more generally demands that researchers working on similar biological questions must be vigilant for non-coding RNA molecules that may not be properly annotated in current genome assemblies.

## Results

### The *ivory* lncRNA promoter is conserved in butterflies and is bound by color pattern factors

As described above, efforts to characterize the function of the *cortex* gene in wing pattern development have proven challenging. We thus sought to identify previously unrecognized features of the locus through a more careful annotation of transcribed elements and chromatin marks within the *cortex* topologically associated domain (TAD) (Fig. 1A). We generated long-read PacBio transcript sequences from a collection of tissues that included wings at different stages of development (Table S1). These data identified a lncRNA that partially overlaps with the 5’ end of the *cortex* gene. The 3’ region of this predicted lncRNA also overlaps with the previously described 78kb *ivory* mutation – a spontaneous deletion identified in *Heliconius melpomene* that produced largely unpigmented butterflies (14) – thus, this novel lncRNA is dubbed *ivory* after the original *Heliconius* mutation.

**Figure 1.**
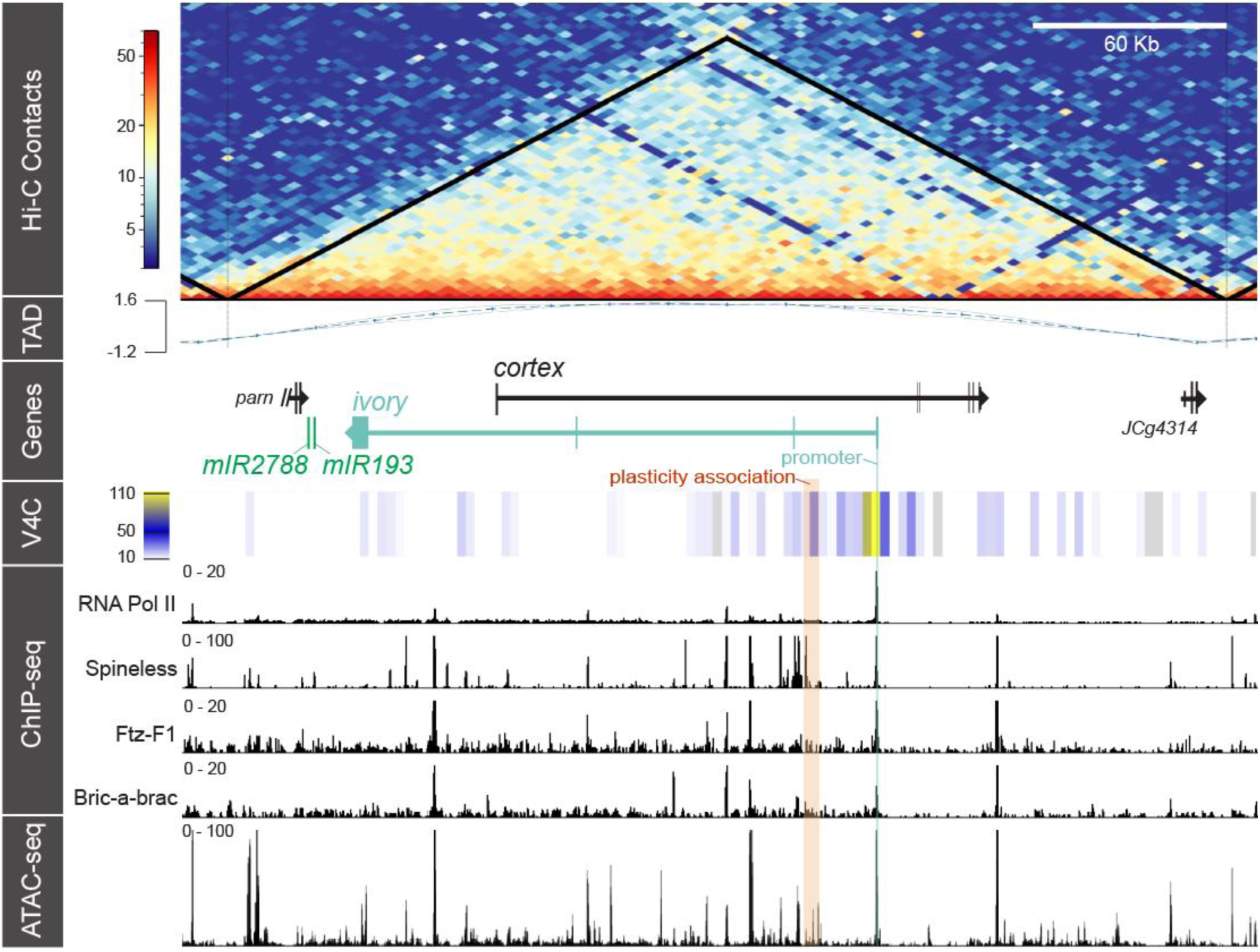
Hi-C chromatin conformation capture reveals a topologically associating domain (TAD) that centers on the overlapping *ivory* and *cortex* genes. The Hi-C Contact heat map displays chromatin interaction strength. The TAD track presents the TAD separation score. The virtual 4C (V4C) track illustrates relative strengths of chromatin contacts specifically with the ivory promoter (yellow). ChIP-Seq tracks annotate binding of RNA Pol II and three transcription factors during early pupal wing development. The orange column highlights a region of nucleotide variant associations linked with natural variation in seasonal polyphenism of *J. coenia* wing coloration (2). The ATAC-seq track illustrates regions of chromatin that are accessible during early pupal wing development.

We next annotated the *ivory* region with RNA Polymerase II (Pol II) ChIP-seq data to identify potential transcription start sites (promoters) within the TAD (Fig. 1). We identified a major Pol II binding site during pupal development, which occurred at the 5’ end of the first exon of *ivory*. To identify CREs of potential interest, we generated and/or compiled pupal wing ChIP-seq data from several transcription factors implicated in pigment regulation (Table S2). We looked at previous ChIP-seq data for the wing pattern transcription factor Spineless (15), which produces similar wing depigmentation mutant phenotypes as *ivory* (16), and generated new ChIP-seq data for Fushi tarazu-F1 (Ftz-F1) and Bric-a-brac (Bab) (Fig. 1), for which mutants have globally reduced or enhanced pigmentation, respectively (Fig. S1). All three transcription factors showed strong binding signals at the presumptive *ivory* promoter, as well at a handful of other apparent CREs within the TAD, as inferred from ATAC-seq data (Fig. 1). Spineless, in particular, showed a strong cluster of binding sites in the first intron of *ivory*, in an interval previously identified as showing nucleotide polymorphism associations with seasonal polyphenism in *J. coenia* (2). A broad comparison with ATAC-seq-annotated genomes of multiple nymphalid butterflies showed that the first exon of *ivory* is conserved across the family, along with many conserved CREs inferred from chromatin accessibility profiles (Fig. S2, Table S3). Together, these data demonstrate activity of the conserved *ivory* lncRNA promoter and transcript during pupal wing development, and that *ivory* is directly bound by color pattern transcription factors Spineless, Ftz-F1, and Bric-a-brac, including Spineless binding at CREs associated with seasonal polyphenism.

### *ivory* plays roles in wing pigmentation and eyespot development

The above data, in the context of previous mapping studies from many different butterfly and moth species, identify *ivory* lncRNA as a strong candidate gene for wing pattern adaptation. This lncRNA escaped annotation in previous mRNA-seq studies, however. Our Iso-Seq data allowed us to identify a long *ivory* transcript, for which our Pol II data imply transcription occurs in developing pupal wings. To assess a potential role for *ivory* in color pattern development we generated two mutant deletion lines using CRISPR/Cas9. We targeted the promoter and first exon of ivory with three single guide RNAs (sgRNAs) (Fig. 2A, Table S4). After intercrossing mosaic knockout (mKO) parents, progeny with strong ventral wing pigmentation phenotypes were selected and crossed. We generated two lines: one with an 18bp deletion (*ivory^Δ18bp^*) of the core promoter that does not affect the first exon of ivory (Fig. 2A), and one with a 17kb deletion (*ivory^Δ17kb^*) that spans the promoter and first exon, but that does not affect any *cortex* coding regions (Fig. 2B). The *ivory^Δ18bp^* lines presented an overall faded phenotype, but still appeared to have most pigment types represented (Fig. 2C). This mutation shows that even minor modifications to the *ivory* promoter are sufficient to produce global quantitative effects on pigmentation.

**Figure 2.**
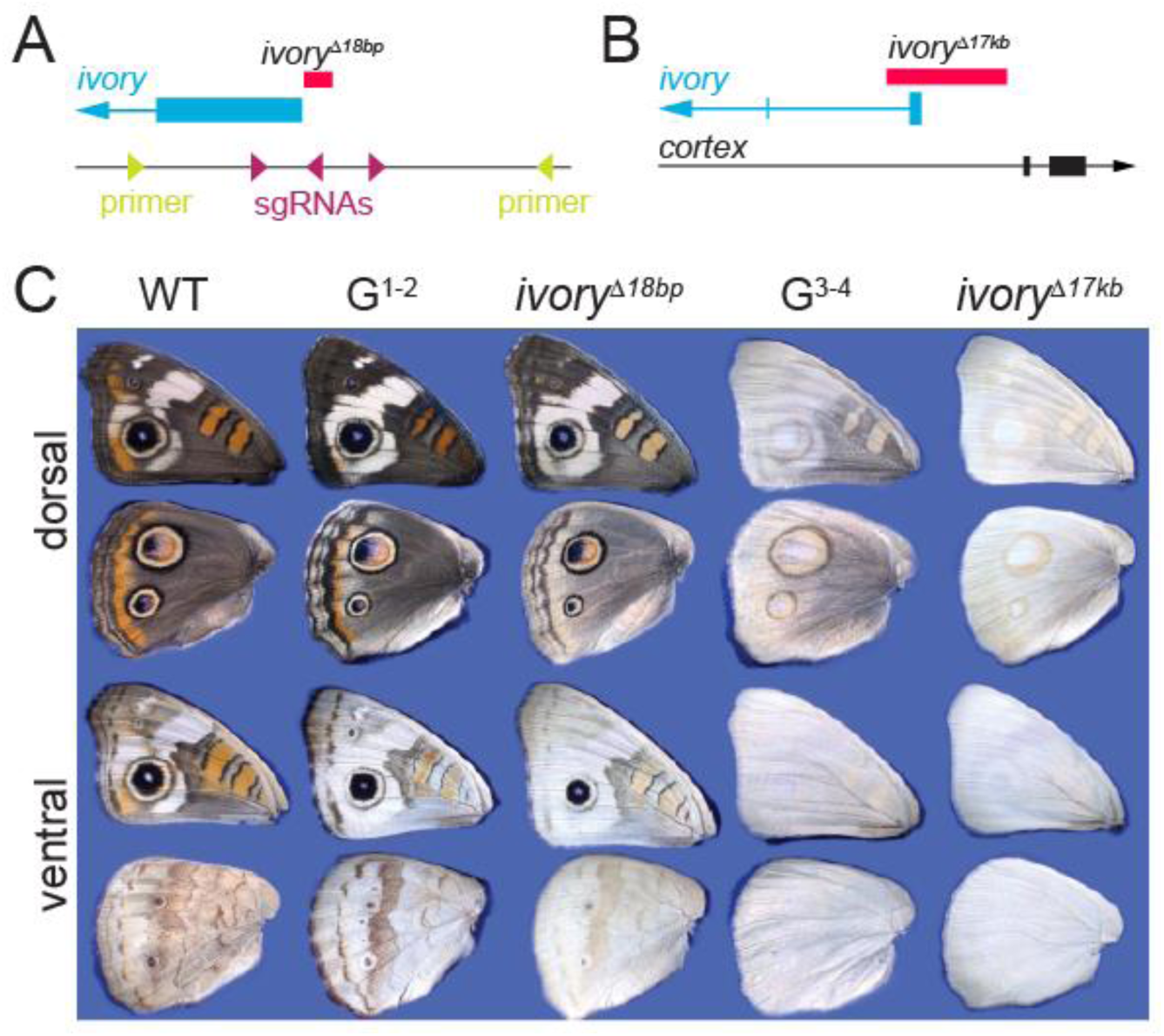
*ivory* promoter deletion lines show reduction and loss of pigmentation. (A) The *ivory^Δp18bp^* deletion (red) is adjacent to the *ivory* 5’ exon, in the promoter. Location of PCR genotyping primers (green) and sgRNAs (magenta) is shown. (B) Expanded view of the *ivory* 5’ region illustrating the location of the *ivory^Δ17kb^* deletion. (C) Phenotypes from the promoter deletion line purification process. In addition to wildtype and homozygous *ivory^Δp18bp^* and *ivory^Δ17kb^* individuals, we also show examples of presumptive heterozygotes from generations 1-2 (G^1-2^) and 3-4 (G^3-4^) of the selection process that illustrate the range of variation we observed. Of note is the quantitative variation in overall pigmentation, but consistent presentation of the melanic center rings of the large eyespots.

The *ivory^Δ17kb^* line, which was confirmed through whole genome sequencing, produced nearly pigmentless butterflies (Fig. 2C). This phenotype resembled the original *H. melpomene ivory* spontaneous mutants (14), where almost all melanins appeared to be lost, and only weak traces of presumptive ommochrome pigmentation remained. *ivory^Δ17kb^* did not prove to be embryonic lethal, although homozygous butterflies were flightless and not as active as heterozygotes or wild types. Heterozygous *ivory^Δ17kb^* mutants showed an intermediate phenotype of partially faded pigmentation. We were able to maintain this line for several generations by occasional backcrossing to the wild type line and genotyping.

One notable observation when comparing the deletion lines was how *ivory^Δ18bp^* individuals showed a faded pigmentation phenotype typical of various *ivory* knockouts, yet retained fully pigmented melanic center rings in the major eyespots that did not appear to show similar fading (Fig. 2C). This contrasts with the *ivory^Δ17kb^* homozygotes which show a complete loss of melanic inner eyespot rings (Fig. 2C). These results suggest that the pigment fading effect can be decoupled from *ivory*’s role in determining internal melanic eyespot rings, possibly through the effects of specific *cis*-elements. Importantly, none of the mutants we observed appeared to show any change in the size or location of identifiable color pattern elements themselves, leading us to infer that the main function of *ivory* lies in pigment regulation, and not pattern formation *per se*.

### *cis*-Regulatory deletions replicate alternate seasonal phenotypes

*J. coenia* presents different ventral hindwing phenotypes depending on seasonal conditions (Fig. 3A). As described above, a region of open chromatin and Spineless binding in the first intron of *ivory* has been genetically linked to variation in this polyphenic response (Fig. 1A, Fig. 3B) (2). To determine how this and other *ivory cis*-regulatory regions may be involved in color pattern regulation, we used CRISPR/Cas9 to generate “shotgun deletion” somatic mosaic knockouts (mKO) at five presumptive *ivory* regulatory regions: the promoter, one upstream element (E149), and three CREs in the polyphenism-associated interval (E205, E209, and E230) (Fig. 3B, Table S5). As previously described, this CRISPR/Cas9 approach targets a CRE with multiple sgRNAs to induce a spectrum of mutations of differing lengths, thus generating G^0^ somatic mosaics that represent a diversity of mutant alleles that allow us to survey the breadth of effects that regulatory mutations can produce (8).

**Figure 3.**
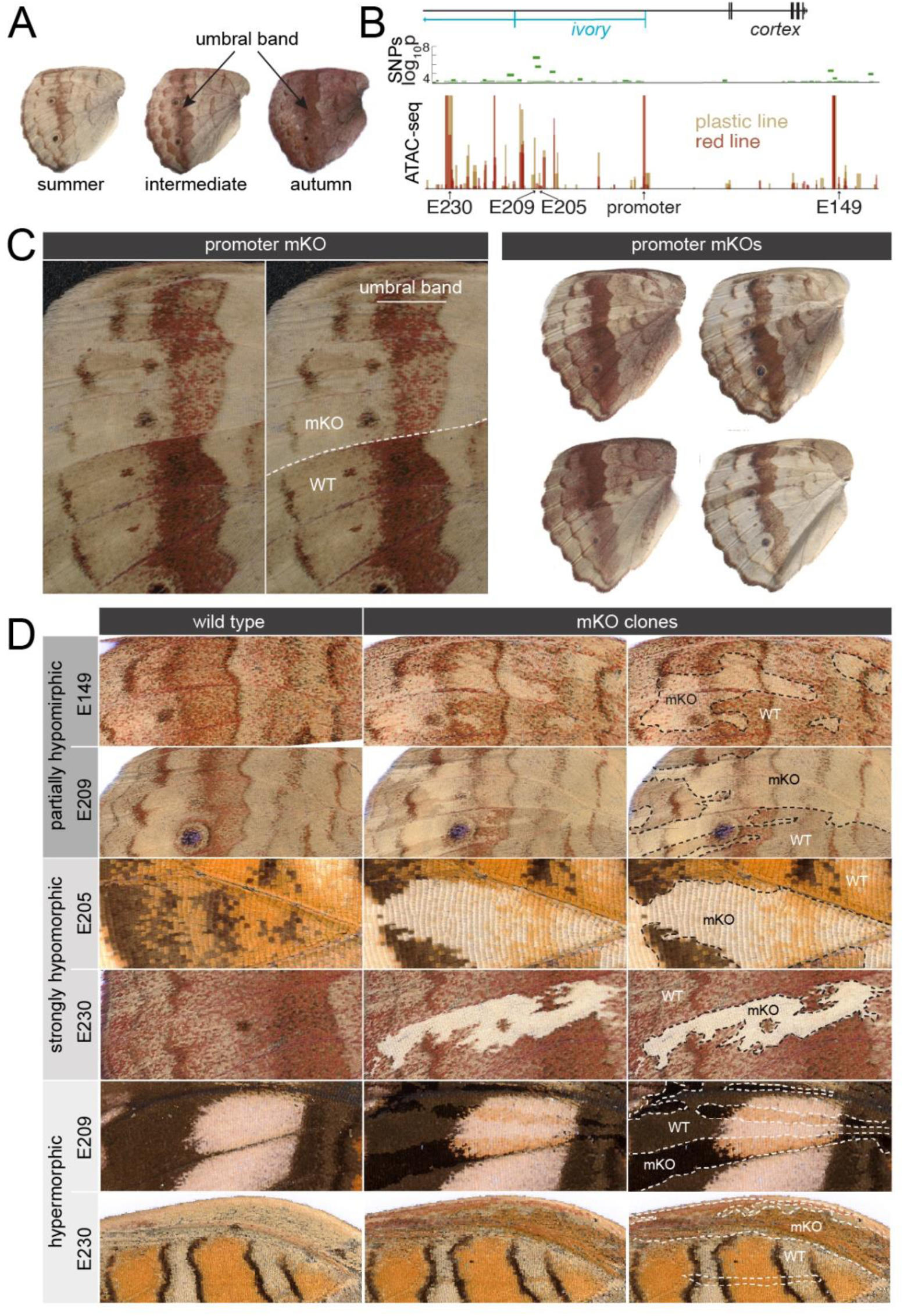
CRISPR mKOs of *cis*-regulatory regions result in several distinct phenotypes. (A) Examples of natural seasonal ventral hindwing phenotypes, including an intermediate form. (B) Locations of single nucleotide polymorphisms (SNPs) linked to variation in seasonal plasticity, along with ATAC-seq peaks from previously published “plastic” and “red” selection lines (2). Targets of CRISPR-Cas9 mKOs are annotated relative to SNPs and ATAC-seq. (C) Examples of promoter mKOs showing partially hypomorphic phenotypes resembling seasonal variation. (D) Examples of the different classes of mKO phenotypes from deletions in the targeted CRE regions.

This exercise produced three distinct phenotypes (Fig. 3C, D, Figs. S3-S7). The first, and most compelling, class of phenotypes was the replication of seasonally plastic color patterns that originally led to the mapping of this locus in *J. coenia*. Specifically, some deletions at all elements generated mosaic clones with partially hypomorphic reduction of dark red pigmentation on the ventral hindwing in a fashion that phenocopied summer phenotypes of *J. coenia* (Fig. 3C, D). In these mutants the “background” beige and brown coloration was minimally affected, while red coloration interspersed across an otherwise beige background field was reduced. Many of these mutants showed reduced or undetectable effects on the dark red umbral band as well (e.g., Fig. 3C), in a manner highly reminiscent of natural seasonal variation (Fig. 3A). Thus, mutant butterflies that would otherwise show an autumn phenotype display mutant clones that resemble the summer phenotype. This contrasted with *ivory* exonic knockouts (e.g. *ivory^Δ17kb^*), or strong CRE hypomorphs, as described below, which showed a uniform loss of almost all pigmentation across the ventral hindwing, including the umbral band (Fig. 3D).

Second, deletions at all elements were all capable of producing strongly hypomorphic mutant clones that resembled the *ivory^Δ17kb^* line, and as evidenced in other species (6, 14) – an almost complete loss of most pigmentation on both dorsal and ventral wing surfaces (Fig. 3D). We infer that these elements function as enhancers, or positive regulators, of *ivory* transcription, and that the fading phenotypes are likely a result of decrease in *ivory* transcription or dosage. The final class of phenotypes, seen in a subset of E209 and E230 mutants, consisted of pigmentation hypermorphs that showed darkening of both dorsal and ventral pattern elements (Fig. 3D). We infer that these CREs possess some silencer-like functionality, and that their deletion likely results in increased *ivory* dosage.

Together, these results demonstrate that *ivory* can be regulated by non-coding sequences in a relatively refined fashion, likely through dosage-like effects on global pigment intensity, but also showing evidence of potentially pattern-specific strength-of-effect (e.g., umbral band and central eyespot rings). Importantly, our experimental replication of nuanced seasonal phenotypes serves to validate previous genetic mapping work that links this locus to color pattern plasticity.

### *ivory* expression is correlated with color pattern

The results above demonstrate roles for *ivory* in pigment pattern development that can be both broad (e.g. overall pigment intensity) and specific (e.g. melanic eyespot rings and seasonal umbral bands). An implication of these results is that this gene has a regulatory program that encodes some degree of pattern- and season-specific activity. To assess this we characterized spatial expression of *ivory* using hybridization chain reaction *in situ* hybridizations in early pupal wings. At this early stage of pupal development we observed high expression levels of *ivory* lncRNAs specifically in melanic color pattern elements, particularly the outer boundaries of the discal bands (Fig. 4A) the eyespot central and outer rings (Fig. 4A, B). This strong association with melanic color pattern elements was consistent with chromogenic ivory *in situs* in other butterflies, as shown in Livraghi et al. (9), and suggests a positive role for *ivory* in regulating these patterns. These expression domains also predict the color patterns that were retained in the *ivory^Δ18bp^* yet lost in the *ivory^Δ17kb^* line.

**Figure 4.**
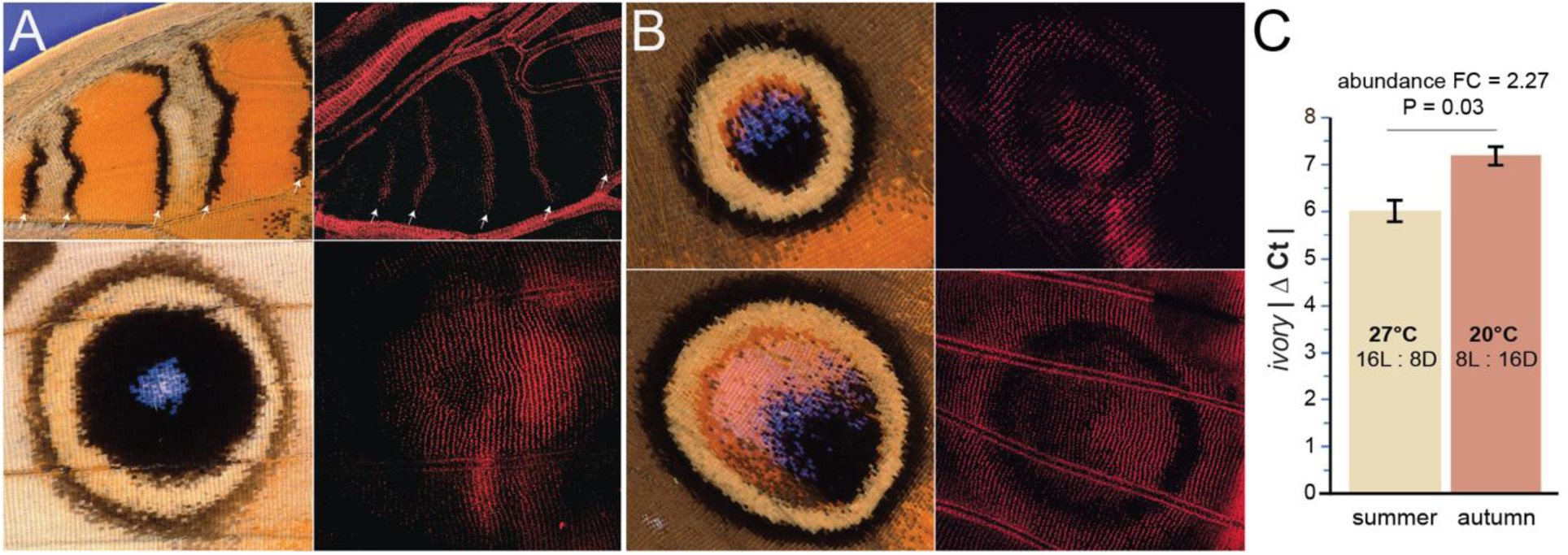
*ivory* expression associated with color pattern phenotypes. HCR *in situ* hybridizations of early pupal wings reveals *ivory* transcripts in (A) ventral forewing color patterns, including black borders of ventral discal bands (white arrows) and the black center ring of large eyespot, and (B) center and rings of dorsal hindwing anterior (top) and posterior (bottom) eyespots. (C) qPCR of mid-stage pupal wings showed 2.27-fold higher *ivory* transcript abundance in wings of butterflies reared under autumn conditions relative to summer conditions (Welch’s *t*-test: P = 0.03, t = 3.88, df = 3; ΔΔCt = -1.18; Rq = 2.27). See Table S6 for details.

Interestingly, at the early stage of development we used for *in situ* hybridization we did not detect *ivory* expression associated with non-black pattern elements such as the umbral band. We speculated that *ivory* expression affecting non-black patterns may occur later in development, as is the case with many pigment-related wing patterning genes (17). We thus used quantitative reverse-transcription PCR to assay *ivory* transcript levels at a stage halfway through pupal development when scales are cuticularizing, just before pigments begin to develop. We surveyed *ivory* transcript levels in wild type butterflies reared under summer conditions (tan phenotype, Fig. 3A), in wild type butterflies reared under autumn conditions (red phenotype, Fig. 3A), and also in the *ivory^Δ17kb^* promoter knockout line with highly reduced overall pigmentation (Fig. 2C), We found that wings from butterflies reared under autumn conditions had *ivory* expression levels more than twice that of butterflies reared under summer conditions (Fig. 4C, Table S6), while *ivory* transcripts were undetectable in *ivory^Δ17kb^* wings (Table S6). Together, these results show that *ivory* transcript levels are strongly influenced by environmental conditions, and that higher levels of *ivory* transcripts are associated with darker pigmentation.

### *ivory* regulates a large and dynamic cadre of transcripts during wing development

The diverse effects of *ivory* on wing coloration, including regulation of multiple pigment types, implies that this lncRNA directly or indirectly regulates multiple biological processes. To assess the transcriptional impact of *ivory* loss of function, we characterized the wing transcriptome at three stages of development in the *ivory^Δ17kb^* line: (1) last (5th) instar wing imaginal discs, when pattern formation begins; (2) pupal wings two days after pupation when patterns are finalized, but before scales mature, and when our *in situ* hybridizations in Fig. 4 were performed; and (3) late pupal wings, 6-7 days after pupation, when scales are nearly mature and pigments are being synthesized. We compared wild type and mutant transcriptomes and identified the most differentially expressed genes at each timepoint (Tables S7-S9). There was an overall large effect of the *ivory* knockout on gene expression during wing development with thousands of significantly differentially expressed transcripts (5^th^: 518; Day 2: 3629; Day 6: 469) at FDR of *p*^adj^ =0.01 (Fig. 5).

**Figure 5.**
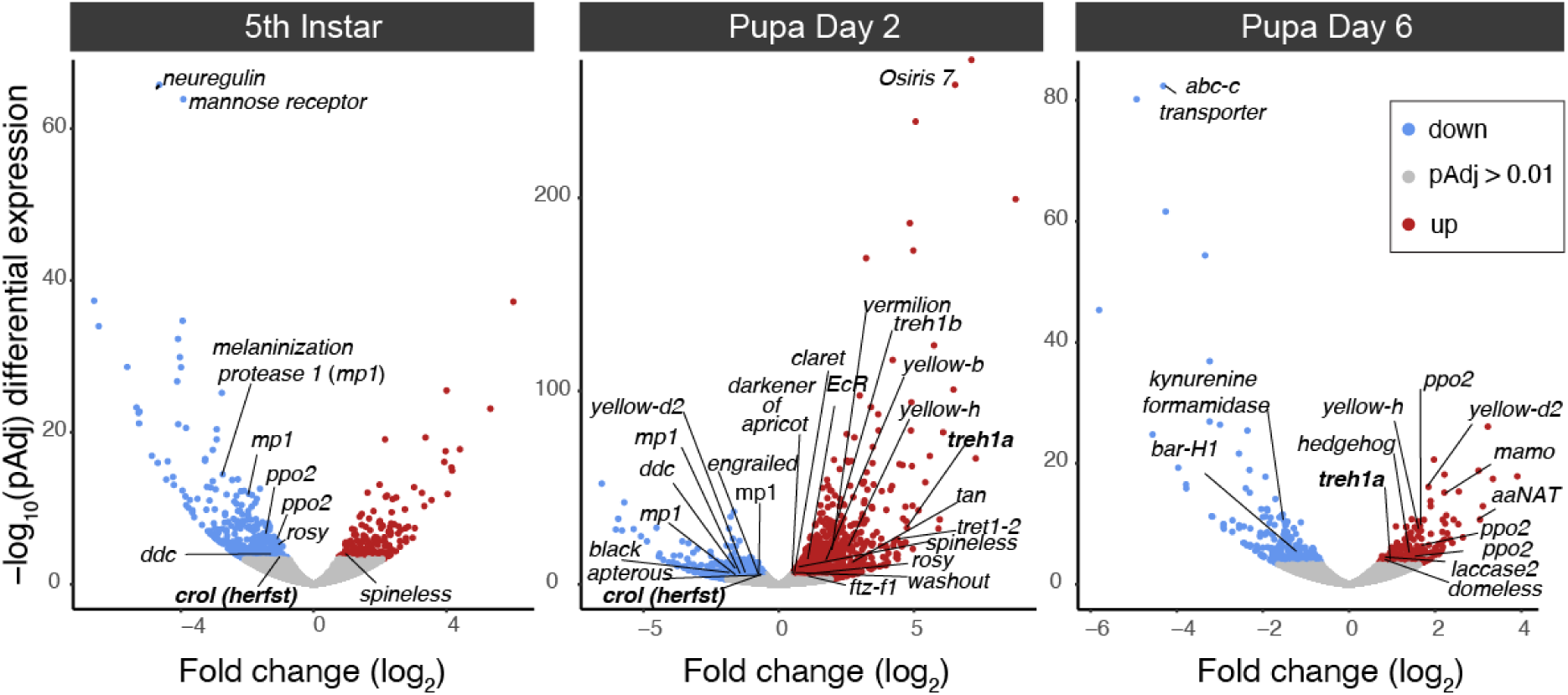
*ivory* loss-of-function causes major changes in gene expression during wing development. Volcano plots illustrate differences in transcript abundance in wild type versus pigmentless *ivory^Δ17kb^* wings at three stages of development. Blue dots mark genes downregulated in mutants and red dots mark genes upregulated in mutants. Genes potentially implicated in pigmentation or wing pattern development are highlighted. Genes previously identified in seasonal plasticity are in bold. *ddc*: *dopa-decarboxylase*; *EcR*: *ecdysone receptor*; *ppo*: *prophenoloxidase*.

We looked specifically at differentially expressed genes with known or suspected roles in wing pigmentation and/or patterning (Table S10) to ask if the *ivory* knockout phenotype is consistent with functions of these genes. In this respect, there were a number of pigmentation genes that were downregulated in knockouts, including the ommochrome gene *kynurenine formamidase*, and melanin genes including *laccase2*, *dopa decarboxylase*, *black*, and *yellow* gene family members *yellow-d2* and *yellow-c* (17). *kynurenine formamidase* was of particular interest because of its high normalized read counts in late pupal wild type wings (> 2k), implying very high expression, perhaps consistent with the whole-wing effects of the ivory loss of function phenotype. Interestingly, there were also a few pigment-related genes upregulated in mutants, including the melanin-related genes *dopamine N-acetyltransfrease* (*Dat*), *mamo*, *tan*, and multiple *prophenoloxidase*- and *yellow*-family genes. *trehalase* and several homologs of *tret 1-2*, a predicted trehalose transporter, were also highly upregulated in early pupal wings, which is notable because *trehalase* knockouts present a summer-morph phenotype similar to *ivory* CRE knockouts (2). A >10-fold change increase in *trehalase* and *tret 1-2* expression in early pupal wings of *ivory^Δ17kb^* thus provides extra support for a functional connection between these loci in the context of wing pigmentation, yet the inverse correlation between their expression suggests a more nuanced relationship than previously surmised.

In terms of known color pattern genes that were differentially expressed, the best supported candidates included *spineless*, *engrailed*, *washout/domeless*, *herfst/crooked legs*, *bar-H1*, and *ecdysone receptor* (2, 16, 18, 19). While it is perhaps premature to speculate about the functional connection between *ivory* and most of these genes, *spineless* stands out for three reasons: [1] knocking it out produces global, wing-wide effects on pigmentation just as *ivory* does; [2] it binds to the *ivory* promoter (Fig. 1), suggesting reciprocal regulation; and [3] it has surprisingly high expression levels in mutant wings during early pupal development, with normalized read counts exceeding 10k in *ivory^Δ17kb^* wings (Tables S7). *ecdysone receptor* is also an attractive candidate because titers of the steroid hormone ecdysone determine seasonal wing pattern phenotypes (2), thus providing a possible mechanism for *ivory*’s role in color pattern polyphenism.

A final notable pattern that emerged from the RNA-seq analysis was the strong upregulation of multiple *Osiris*-family genes in in *ivory^Δ17kb^* wings during early pupal development, in terms of fold change, significance, overall read count magnitude, and overall number of *Osiris* genes (Table S8). This gene family is thought to encode transmembrane transporters, although *Osiris* genes have not been well characterized outside of tracheal function in *Drosophila* (20), and they have not previously been linked to butterfly wing pigmentation. This gene family may provide candidates for future functional assessment

## Discussion

In this study we show that the lncRNA *ivory* regulates *J. coenia* color patterns, and that perturbation of *ivory* CREs can produce nuanced color pattern variations, including replication of seasonal phenotypes (2, 21). In terms of color pattern evolution, these results are of broad interest because they prompt a rewrite of our understanding of the *cortex* locus, which has received significant attention for its association with wing pattern adaptation across numerous species of moths and butterflies. Our findings, particularly in the context of recent *cortex* deletion lines lacking color pattern phenotypes (9), show that *ivory*, not *cortex*, is the color pattern gene of interest. Therefore, important molecular case studies of industrial melanism (3), Müllerian mimicry (6), and crypsis (4) should perhaps be revisited. Our work provides a cautionary example for anyone working to identify genes underlying variation that we need to look more carefully for evidence of causative non-coding transcripts.

More broadly, *ivory* represents a potentially novel case study of deeply conserved non-coding RNA underlying repeated instances of adaptation. Non-coding RNAs are now known to be abundant in genomes, and are being increasingly recognized as important regulators of gene expression (10). lncRNAs like *ivory* are generally thought to act as *trans*-regulators of gene expression through interactions with various protein, RNA, and DNA targets. This process may include intermediate posttranscriptional processing of lncRNAs into smaller effector RNAs with specific functions (22, 23). Indeed, a preprint posted in parallel with our study presents evidence that a micro-RNA (*miR-193*) immediately 3’ of *ivory* may be derived from the nascent *ivory* transcript to play a role in pigment regulation (24). Thus, it is interesting to speculate whether *miR-193* might be an effector molecule for *ivory* color pattern functionality, perhaps as a *trans*-regulator of pigmentation genes. Pending further work, however, the specific molecular mode of *ivory* action remains an open question.

While lncRNAs have been linked to various human health conditions, including morphological and neurodevelopmental disorders (25, 26), little is known about how they may function in contexts such as adaptation or phenotypic plasticity. One of the few case studies that directly implicates any kind of non-coding RNA in trait adaptation concerns the role of small RNAs at the *YUP* locus in monkeyflowers, which affect carotenoid flower pigmentation (27). It is interesting that the *YUP* case study involves pigmentation, as does our *ivory* study. The similarities end there, however, as the *YUP* locus encodes multiple small RNAs that are evolutionarily labile, and individual RNAs can be specific to individual species. The *ivory* gene appears to behave in an evolutionary mode more consistent with other wing pattern adaptation loci like *optix* (a transcription factor) or *WntA* (a ligand), where a deeply conserved gene with a core developmental role diversifies phenotypes via *cis*-regulatory innovation (8, 28).

Our study was initially prompted by *J. coenia* DNA sequence associations that link a cluster of intronic CREs to variation in seasonal color pattern plasticity (2). Targeted deletions in this regulatory region caused a switch from autumnal color pattern to a summer color pattern, and thus appear to confirm a role for these CREs in modulating seasonal color patterns. What is interesting beyond simply being a validation of previous association work, however, is that these phenotypes also demonstrate that CREs of *ivory* can exercise nuanced, quantitative control over the color identity of large fields of cells. Natural butterfly color pattern diversity clearly consists of much more than binary cell fate decisions across a fixed palette of colors - there is also natural quantitative variation in hue, intensity, and semi-stochastic expression of local color schemes across specific wing fields. Thus far, however, we know very little about the developmental genetic basis of variation in these kinds of quantitative color qualities (29). *ivory* stands out for playing this kind of role, and we wonder whether its identity as a lncRNA may be tied to this nuanced functionality in color regulation.

lncRNAs are known for their gene regulatory functions (25), but they can also play other functional roles within cells, including in protein interactions, organelle regulation, etc. (10). We were thus curious to determine to what extent differential expression of *ivory* affected gene expression. Does it regulate many genes, or only a handful of key pigmentation genes? Or perhaps it has no effect on gene regulation, and plays a direct role in pigment synthesis, precursor transport, or other aspects of scale cell biology. Our developmental RNA-seq data from wings of *ivory^Δ17kb^* individuals, with nearly pigmentless *ivory* loss-of-function phenotypes, show clearly that *ivory* has strong effects on transcript abundance during wing development, and that many genes show evidence of both upregulation and downregulation in mutant wings (Fig. 4). The transcripts include a number of candidate pigmentation and wing patterning genes that are suggestive for explaining the pigment-related phenotypes (Table S10). Downregulation of various melanin and ommochrome genes (e.g. *dopa decarboxylase*, *black*, *kynurenine formamidase*) in *ivory* mutants makes immediate sense because of the pigment loss phenotypes, but also compelling is the upregulation of other pigmentation genes, such as *mamo*, a recently characterized melanin repressor in the silkmoth (30). Of course, however, we do not know which genes or transcripts are direct targets of *ivory* activity versus indirect downstream effectors. Livraghi et al. (9) speculate that a primary function of *ivory* function lies in regulating melanin synthesis genes, which we agree is likely to be true. But our RNA-seq data also make it clear that *ivory* has a strong and broad influence over the expression of many genes, even in larval imaginal discs, which may help explain its apparent effects on other non-melanic pigment types as well. Future work on this lncRNA would benefit from characterizing its molecular mode of action and identifying its direct targets during wing development. Through more functional work we will be able to understand how lncRNAs like *ivory* facilitate adaptive variation and plasticity in rapidly evolving gene regulatory networks.

## Materials and Methods

### Butterfly husbandry

Our primary *J. coenia* stock originated from Durham, North Carolina, and was reared in the lab at 27°C, 16 hrs light: 8 hrs dark on an artificial wheat germ diet supplemented with dried *Plantago lanceolata*, as previously described (31). We used a genetic assimilation “red line” of this stock with constitutive autumn red ventral hindwing coloration (2) to enhance detection of CRISPR/Cas9 mutation effects on red pigment patterns. Genetic mutant lines were visually assessed by pigment fading in the ventral wing (Fig. 2C).

### Iso-Seq long read transcriptome annotation

For whole organism long read RNA annotations we sampled an array of tissues at different developmental stages (Table S1). Tissues were dissected into Trizol and stored at -80 °C. RNA was extracted using RNeasy Mini Kit (Qiagen) with on column DNase treatment and quantified with a Qubit RNA BR Assay kit (Molecular Probes). RNA samples were sent to Maryland Genomics (University of Maryland School of Medicine) for quality control (RIN = 9.9), full length cDNA synthesis and Iso-Seq library preparation. Libraries were sequenced on a PacBio Sequel II 8M SMRT Cells on a 30 hr movie run. The sequencing run provided 1.1 TB of data that was concatenated using the PacBio SMRT Iso-Seq pipeline (https://www.pacb.com/support/ software-downloads/) to generate a circular consensus sequence file (.ccs) along with high quality (hq) and low quality (lq) full length transcript files. A SMRT PacBio wrapper (PBMM2) based on pairwise alignment (MINIMAP2) script was applied using the .css file against the jcgenv2.fa genome (Lepbase). The generated alignment file (.bam) was used for downstream transcriptome re-annotation using the BRAKER3 pipeline (32). Briefly, BRAKER 3 incorporates transcript selector algorithm TSEBRA with AUGUSTUS predictions through BRAKER1 and BRAKER2 pipelines. This allowed us to combine long read sequences (this manuscript) with previous short read data (2) to produce an updated version 3 *J. coenia* transcriptome annotation (Jcv2_OGSv3.gff). Assessment of completeness was performed with BUSCO v5.4.7 with complete BUSCO score of 97.9 % (Table S11). The v3 *J. coenia* gene predictions are available on the Dryad repository (DOI: 10.5061/dryad.qz612jmpb).

### Chromatin immunoprecipitation and sequencing

Pupal wing tissue for chromatin immunoprecipitation of Bric-a-brac, Ftz-F1, and Pol II bindings sites was collected and processed as previously described (2, 18). Further information on antibodies, pipelines, and dataset sources used for this study are found in Table S2. ChIP-seq and input libraries were prepared using an NEBNext DNA Ultra II kit without size selection. Libraries were sequenced using the Cornell Biotechnology Resource Center Genomics Facility on an Illumina NextSeq 500/550 at 20 million bp reads for treatment (ChIP) and 30 million for control (input).

### Hi-C, ATAC-seq, and ChIP-seq dataset analysis

Hi-C derived TAD boundaries and comparative alignments centered on ATAC-seq peak data within the *cortex* locus (Fig. 1) were adapted from Mazo-Vargas et al. (7). We generated a virtual circular chromosome conformation capture (4C) plot to visualize chromatin interactions centered on the *ivory* promoter as previously described (2, 33). Comparative noncoding genome alignments for Scaffold 15 were performed using HALPER on *J. coenia* wing tissue ATAC-seq peak calls (MACS2) (7). Peak calls representing sites of open chromatin were given a score from (0.001 – 1.000), with the lowest score representing regions of least conservation and a score of 1.0 representing conservation in at least eight of twelve species used in the Mazo-Vargas et al. (7) alignment. Pupal wing ATAC-seq datasets of representative nymphalid clade butterfly species were gathered from previous studies: *J. coenia, Vanessa cardui*, *Heliconius erato demophoon*, and *Danaus plexippus* (Table S3). Peak calls for regions of high ATAC-peak conservation were imported into Geneious Prime (v2023.1.1) and manually annotated for position and chromatin accessibility (i.e. ATAC-seq signal) (Table S12). ATAC-seq peaks below (RPKM normalized count values < 10) were characterized as not accessible and marked with a gray line in Fig 1. ChIP-Seq alignment data for the transcription factor Spineless was taken from van der Burg et al, (12), and we generated new Bric-a-brac (three replicates), Ftz-F1 (two replicates), and Pol II (three replicates) ChIP-seq data as described above. ChIP libraries were aligned to the lepbase.org *J*. *coenia* genome version 2 (jcv2). Alignment, duplicate removal, and peak calling was done as previously described (18).

### CRISPR/Cas9 targeted mutagenesis and genotyping

To generate *ivory* loss-of-function lines we designed two sgRNAs targeting the 5’ promoter where Pol II binds (Table S4). We carried out sgRNA-Cas9 embryo injections in *J. coenia* as previously described (34), with modification to microinjection needle shape pull on Sutter-P97 (Sutter Instruments). We selected G^0^ adults with large clonal mosaics for sib mating, the resulting G^1^s were purified to two lines: one with highly faded pigmentation, and one with a nearly complete loss of pigmentation. Whole genome Illumina sequencing of three individuals from each line confirmed that the former had an 18bp promoter deletion (*ivory^Δ18bp^*), and the latter a 17kb deletion (*ivory^Δ17kb^*) (Fig. 3A, B).

To test the function of *ivory* CREs in red line *J. coenia* we generated somatic mKO mutants using sgRNA pairs targeting selected ATAC-seq peaks, as previously described (8). This approach generates mosaic G^0^ individuals, where bilateral asymmetry of mKO clones provides a useful assay for ruling out natural variation or other non-CRISPR effects that would otherwise produce symmetrical phenotypes (17). Cas9 targeting was confirmed by PCR and sequencing of selected mutants for each locus (Table S5).

### Hybridization chain reaction *in situ* hybridization

The first exon of *ivory* was used by to design hybridization chain reaction probe pairs with sequences unique to the *ivory* transcript (Molecular Instruments, *ivory-B1* probe lot RTG621)). Wings were removed from 2 day old pupae and *in situs* were performed following the protocol of Chatterjee et al. (35). B1 hairpin amplifiers were tagged with AlexaFlour 647 fluorophores, and wings were imaged on a Leica Stellaris 5 confocal microscope.

### qPCR analysis of *ivory* transcript levels

We extracted RNA from whole wing tissue of 48% development pupa (4 days after pupation at 27°C, and 8 days after pupation at 20°C) of *ivory^Δ17kb^* individuals raised under summer conditions (27°C; 16L:8D), wild type individuals raised under summer (27°C; 16L:8D) and autumn conditions (20°C; 8L:16D), respectively, as previously reported (2). We gathered three biological replicates for each stage and extracted total RNA from combined forewing and hindwing tissue using the MagMAX Total RNA isolation kit (Thermo Fisher Scientific). Total RNA was DNase treated and used to synthesize cDNA using the TaqMan Advanced miRNA cDNA Synthesis Kit (Life Technologies) with random primers from High-Capacity cDNA Reverse Transcription Kit (Thermo Fisher Scientific). We performed qPCR using PowerTrack™ SYBR Green Master Mix (Thermo Fisher Scientific) using primers to amplify the first exon of *ivory* (forward 5’-ATTTACGGTCCGCCTGTTTCG; reverse: 5’-CCACAATCACAGTACCACATCAA) and the control housekeeping gene *rps13* (forward 5’-GTAAAGGTATCTCACAGTCGCG; reverse 5’-GATCAGGTAGTACAGATCCTCGG). Cycle threshold (Ct) values for *ivory* were normalized using *rps13* Ct data to generate ΔCt values (Table S6). Experimental rearing treatments were compared using a two-tailed Welch’s *t*-test for significance, and the ΔΔCt method to calculate relative expression fold change (Rq) of autumn relative to summer treatments (36).

### *ivory^Δ17kb^* mRNA sequencing and expression analysis

We dissected wings from late 5^th^ instar larvae, day 2 pupae, and day 6 pupae of red line and *ivory^Δ17kb^ J. coenia.* Each of the three experimental replicates for each stage represented all forewings and hindwings pooled from three individuals and stored and homogenized in Trizol (Thermo Fisher Scientific). PolyA-enriched transcripts were isolated using RNeasy Mini Kit (Qiagen) and bidirectionally sequenced by Novogene Co, Ltd. *ivory^Δ17kb^* mutants were genotyped with TaqPhire Pol Direct-Tissue PCR (Invitrogen), DNA was extracted from a third to fourth instar caterpillar spine and added to a reduced 20 µl lysis reaction mix, 1µl of this lysis reaction was used as template for PCR. RNA-Seq datasets were aligned using HISAT2 on jcgen_v2.fa genome followed by Featurecounts using the Iso-Seq informed v3 annotations (Table S1). Refer to Dryad for predicted amino acid sequences corresponding to Jcv2_OGSv3.gff (DOI: 10.5061/dryad.qz612jmpb). Differential gene expression analysis was performed using default settings for DESeq2 on the Galaxy Server webtools (37).

## Acknowledgments

We thank Samantha Pires and Julia Marie Forte for help with butterfly rearing, and Arnaud Martin and Joseph J. Hanly for helpful comments on the manuscript. This work was funded by U.S. National Science Foundation Grants IOS-2128164 and DEB-2242865 to R.D.R.

---

**Figure S1.**
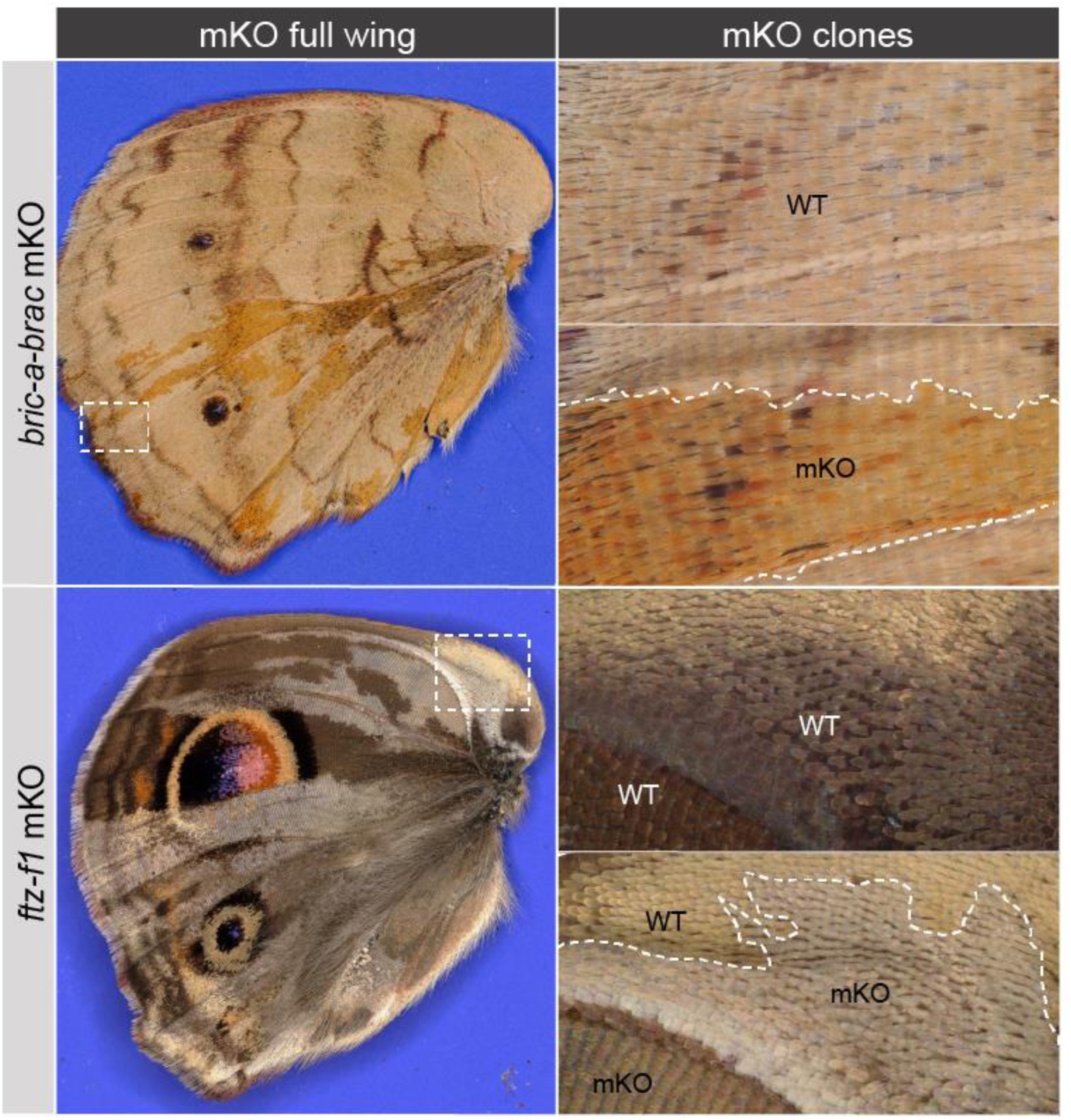
CRISPR deletions of transcription factor genes *bric-a-brac* and *ftz-f1* produce positive and negative global effects on pigmentation, respectively. mKO of *bric-a-bac* shows hyperpigmentation of ommochrome scales in ventral hindwings. mKO of *ftz-f1* results in strong reduction or loss of pigmentation and possible structural effects as well.

**Figure S2.**
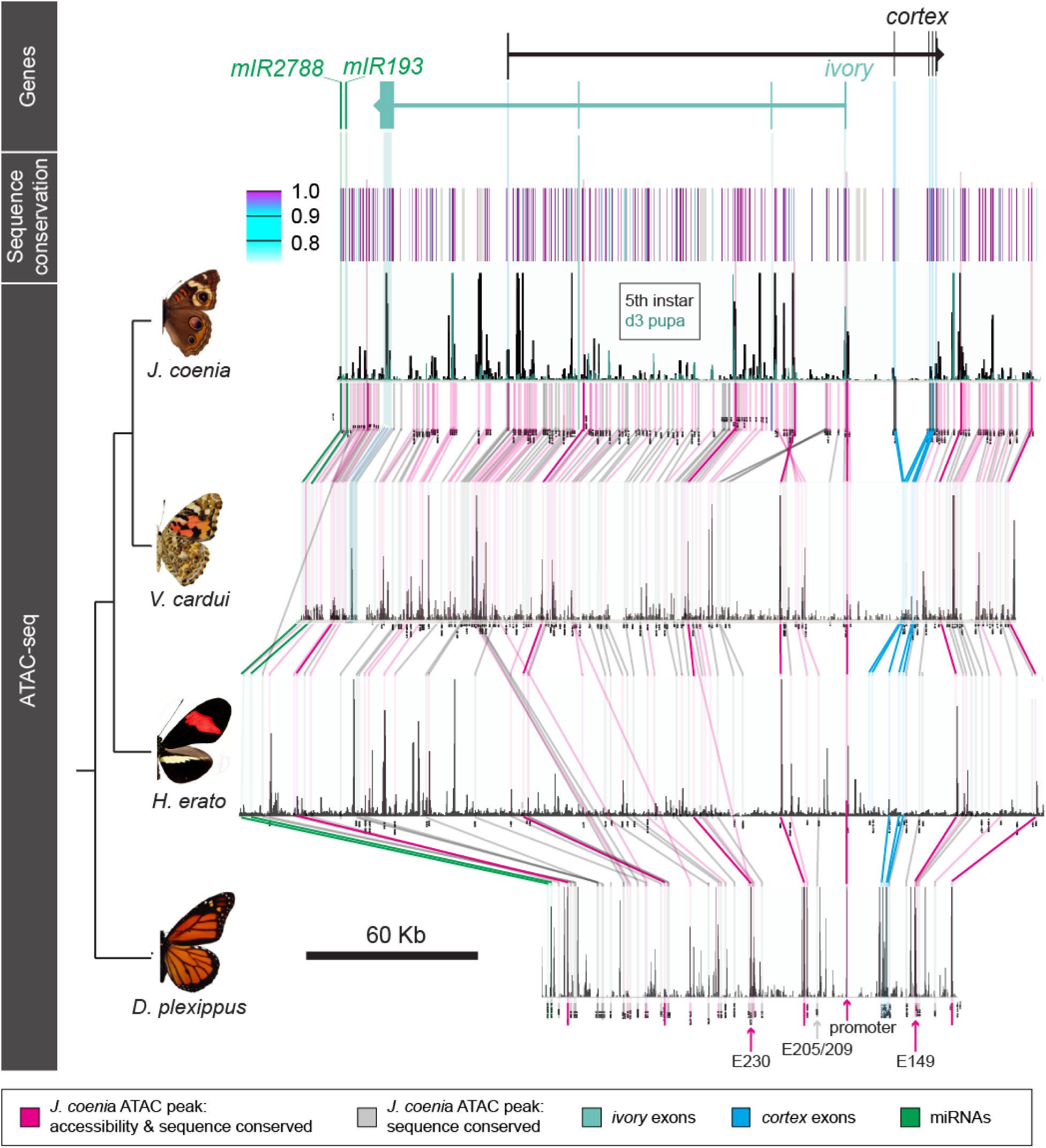
The *ivory / cortex* locus is deeply conserved in nymphalid butterflies. The sequence conservation track shows HALPER sequence similarity scores derived from a comparison between the pictured species. The ATAC-seq tracks present chromatin accessibility data from day 3 pupal wings, with the exception of the *J. coenia* track, which displays both day 3 pupal (green) and 5^th^ instar (black) ATAC-seq data. Vertical bars represent detectable sequence homology between the pictured nymphalid species, for the classes of elements color-coded. The magenta elements, which show conservation in both sequence and accessibility across species, are detailed in Table S12.

**Figure S3.**
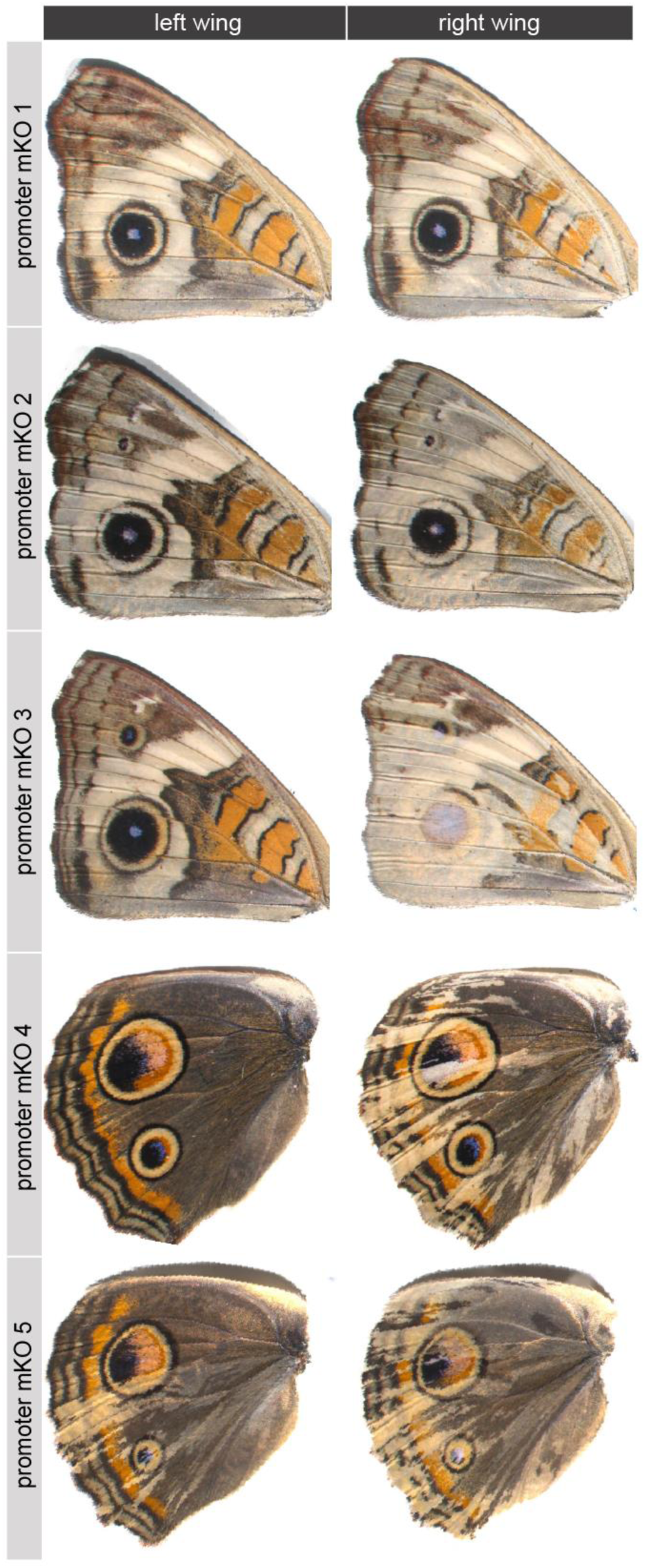
mKOs from sgRNAs targeting the *ivory* promoter (see Fig. 2A). Shown are contralateral pairs of wings from single individuals, emphasizing the asymmetrical mosaicism if G^0^ mKOs.

**Figure S4.**
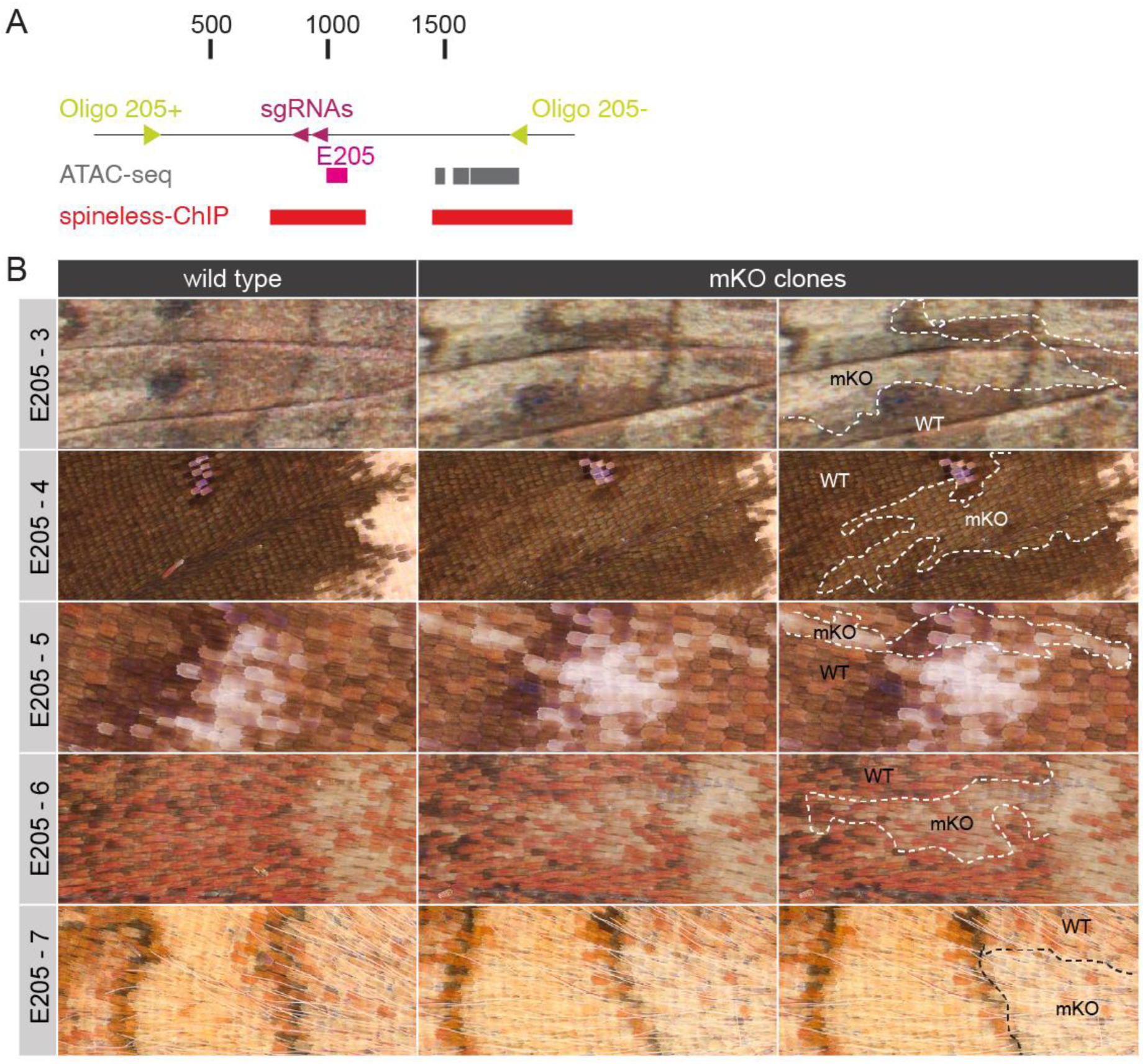
mKO phenotypes from CRISPR deletions of CRE 205. (A) A map of sgRNAs, including ATAC-seq and Spineless ChIP-seq peak calls. (B) mKO individuals showing pigmentation phenotypes.

**Figure S5.**
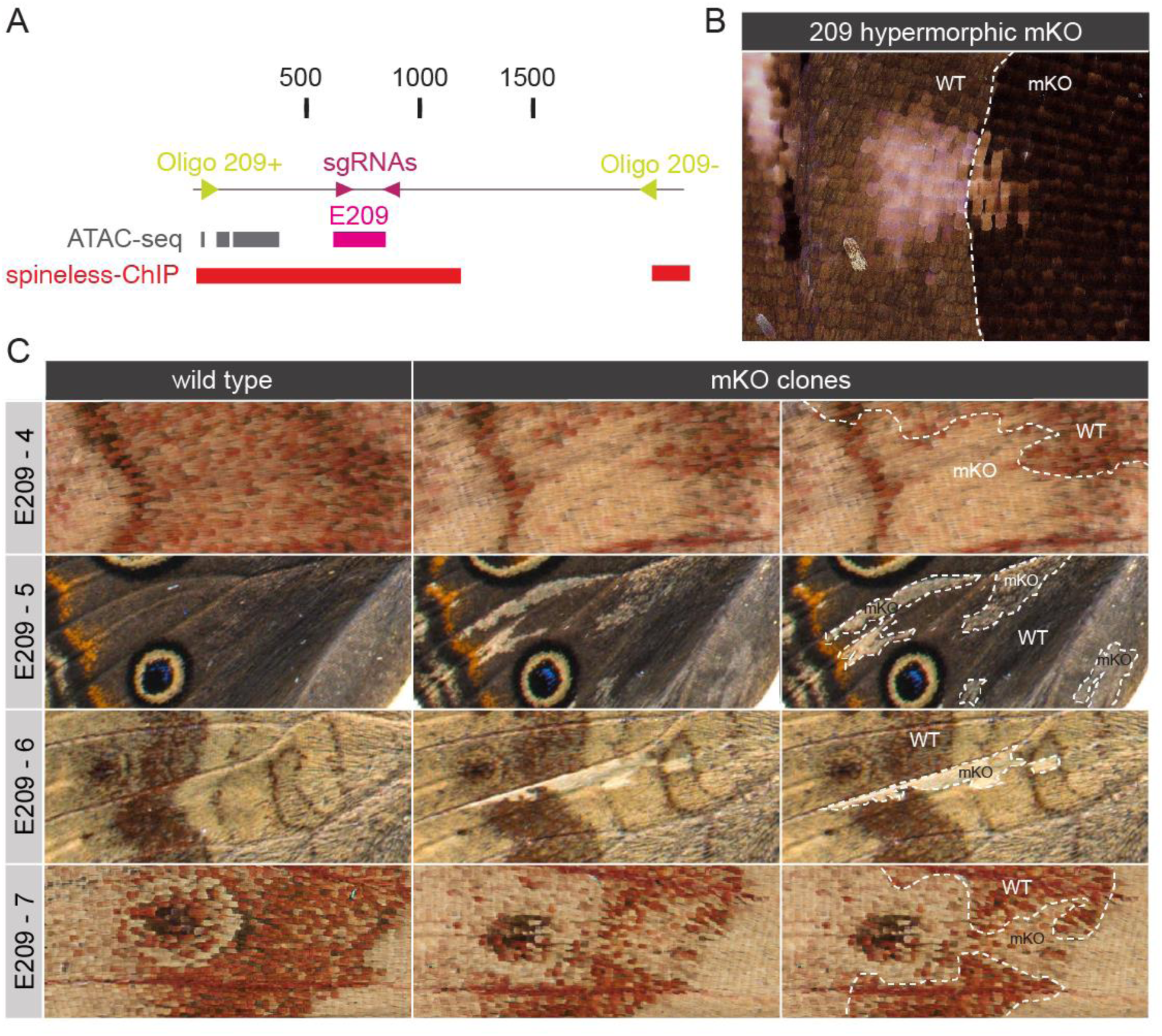
mKO phenotypes from CRISPR deletions of CRE 209. (A) A map of sgRNAs, including ATAC-seq and Spineless ChIP-seq peak calls. (B) A high magnification image of a hypermorphic clone boundary. (C) mKO individuals showing various pigmentation phenotypes.

**Figure S6.**
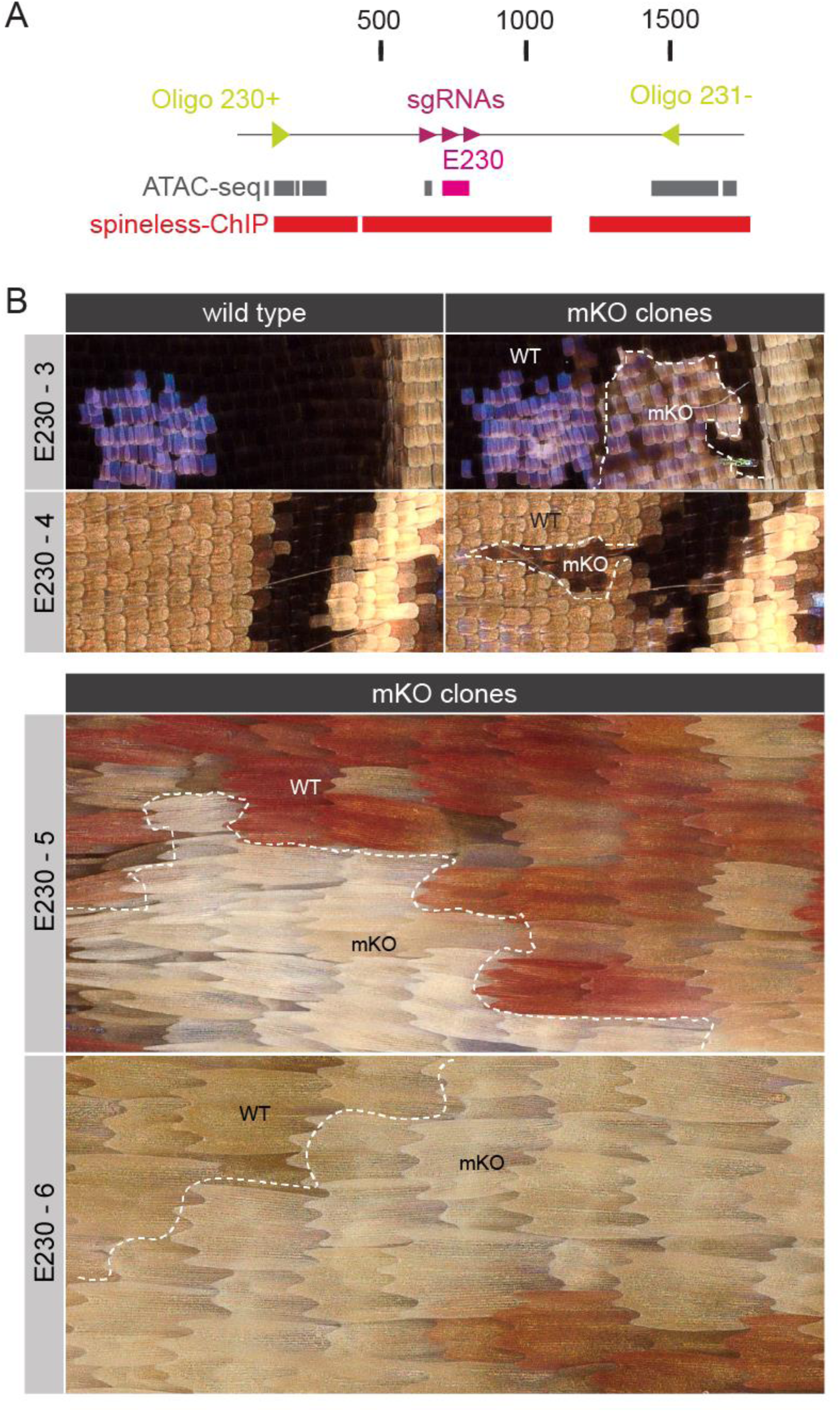
mKO phenotypes from CRISPR deletions of CRE 230. (A) A map of sgRNAs, including ATAC-seq and Spineless ChIP-seq peak calls. (B) mKO individuals showing pigmentation phenotypes.

**Figure S7.**
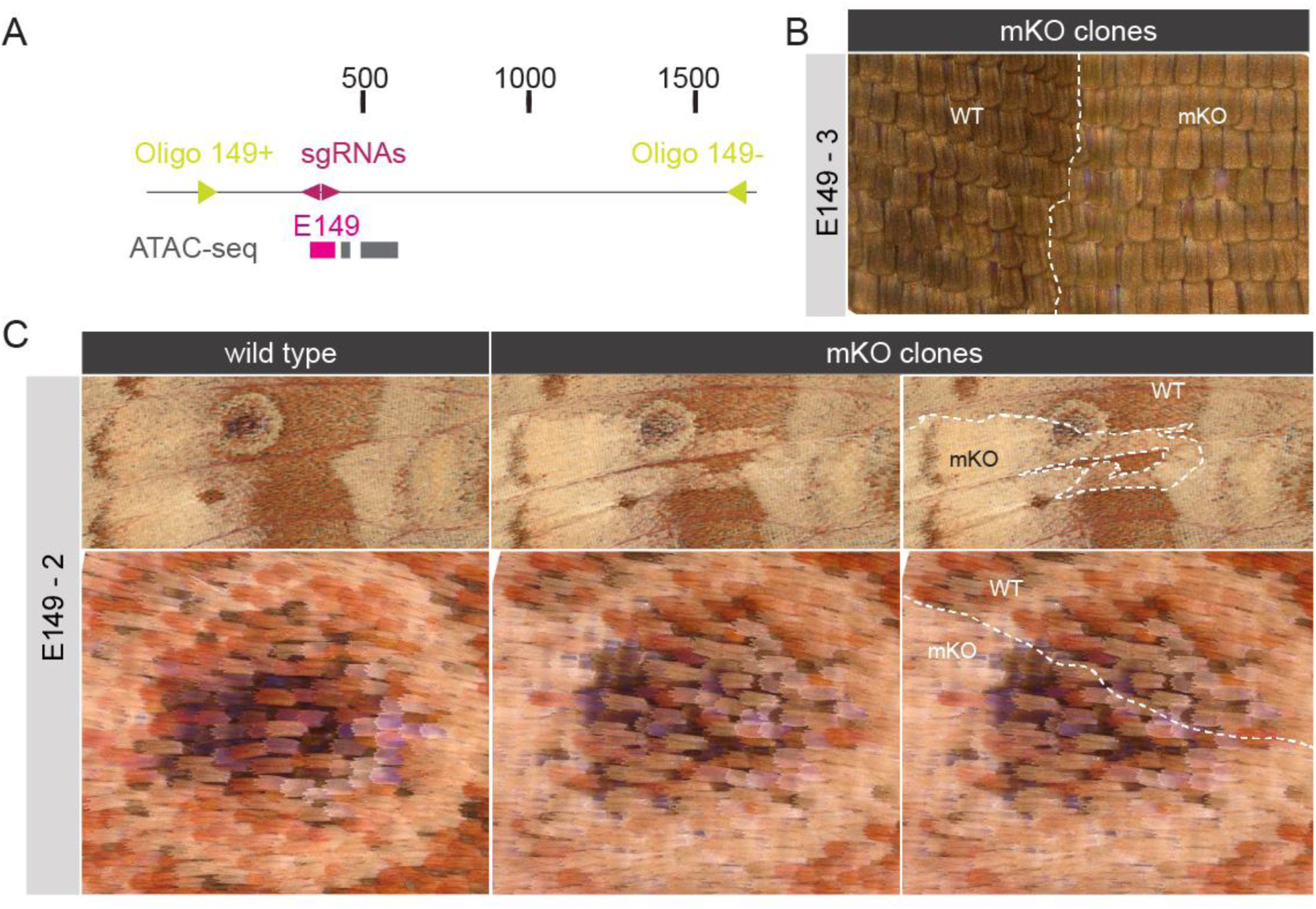
mKOs phenotypes from CRISPR deletions targeting of CRE E149. (A) A map of sgRNAs, including ATAC-seq peak calls. (B) An example mKO individual showing a partially hypomorphic pigmentation phenotype. (C) Another mKO example showing partially hypomorphic clones.

